# Research applications of primary biodiversity databases in the digital age

**DOI:** 10.1101/605071

**Authors:** Joan E. Ball-Damerow, Laura Brenskelle, Narayani Barve, Pamela S. Soltis, Petra Sierwald, Rüdiger Bieler, Raphael LaFrance, Arturo H. Ariño, Robert Guralnick

## Abstract

We are in the midst of unprecedented change—climate shifts and sustained, widespread habitat degradation have led to dramatic declines in biodiversity rivaling historical extinction events. At the same time, new approaches to publishing and integrating previously disconnected data resources promise to help provide the evidence needed for more efficient and effective conservation and management. Stakeholders have invested considerable resources to contribute to online databases of species occurrences and genetic barcodes. However, estimates suggest that only 10% of biocollections are available in digital form. The biocollections community must therefore continue to promote digitization efforts, which in part requires demonstrating compelling applications of the data. Our overarching goal is therefore to determine trends in use of mobilized species occurrence data since 2010, as online systems have grown and now provide over one billion records. To do this, we characterized 501 papers that use openly accessible biodiversity databases. Our standardized tagging protocol was based on key topics of interest, including: database(s) used, taxa addressed, general uses of data, other data types linked to species occurrence data, and data quality issues addressed. We found that the most common uses of online biodiversity databases have been to estimate species distribution and richness, to outline data compilation and publication, and to assist in developing species checklists or describing new species. Only 69% of papers in our dataset addressed one or more aspects of data quality, which is low considering common errors and biases known to exist in opportunistic datasets. Globally, we find that biodiversity databases are still in the initial stages of data compilation. Novel and integrative applications are restricted to certain taxonomic groups and regions with higher numbers of quality records. Continued data digitization, publication, enhancement, and quality control efforts are necessary to make biodiversity science more efficient and relevant in our fast-changing world.

## I. INTRODUCTION

Online databases with detailed information on organism occurrences collectively contain well over one billion records, and the numbers continue to grow. The digitization of natural history specimens (1,2) and development of online platforms for citizen science (3) have driven a steady accumulation of species occurrence records over the past decade. Each data point provides details on the taxonomic identification, date collected or observed, location, and name of the collector or observer for an organism. Applications of these primary biodiversity data are varied—such data have historically helped determine harmful effects of pesticides, document spread of infectious disease and invasive species, monitor environmental change, and much more (4–9). The overall goal of this paper is to quantitatively determine how researchers are using open-access data in published work, focusing on the past decade, when growth of online biodiversity databases has been most rapid. As one illustration of that growth, the Global Biodiversity Information Facility (GBIF) has grown from provisioning just over 200 million records in 2010 to over 1.08 billion records today, a greater than fivefold increase (10).

Museums and funding agencies have invested considerable resources to digitize information from natural history specimens, make their data openly accessible (11,12), and sustain platforms to provide access to those data. Such efforts unlock previously inaccessible data and expand their availability to researchers around the world. However, the task of digitizing highly diverse groups, such as insects, has been particularly difficult. Estimates suggest that only 10% of biocollections worldwide are available in digital form (13), and it would take many decades to completely digitize estimated holdings at current rates (14). While efforts towards workflow optimization will undoubtedly improve efficiency in certain areas (12,15–18), it is critical that the biocollections community prioritize efforts; we must advocate for continued digitization through production of innovative data products, tools, interdisciplinary collaborations, and by highlighting research that requires primary biodiversity data (3,19–21). The greatest returns on digitization investments will result from expanded use of collections data and by linking a wide array of biotic and abiotic data (1,11). Linked data environments are in high demand (22,23), are growing rapidly, and provide the greatest potential for data discovery and use (1).

The biggest obstacle for biodiversity data users is obtaining records of sufficient quantity and quality for the region and taxonomic group of interest (23,24). Many taxa and regions are still highly under-sampled or completely unrepresented (e.g. rare taxa, regions that are difficult to access) in online databases (25–27), particularly for less known and highly diverse invertebrates (28,29). When data are available, they must be checked for common errors and biases known to occur in opportunistic datasets that are often assembled over long time periods (e.g. 30)—a task that is labor-intensive (31). Species identity and locality are the most error-prone aspects of collection information (7). Estimates for rates of collection misidentification range from 5-60% (11,32,33), but if specimens exist, this information can be verified or corrected by taxonomic experts. Specimen images, while not always useful for diagnosis, can often help—particularly when they meet the criteria for taxonomic-grade imaging. Even with correct identification, names in species occurrence repositories may still be incorrect and need validation (34). For many broad-scale studies, erroneous records primarily lead to overestimation of species richness in areas outside centers of diversity (31). Geographic errors may be more readily corrected and associated with appropriate uncertainty estimates using standardized methods (35) and online tools (i.e. GEOLocate, www.geo-locate.org). Digitization of species occurrence records makes it easier to identify questionable records by providing quick access to data and identifying outliers. Further, data services are becoming more sophisticated in automatically addressing some data quality issues (36,37). However, it is possible that many studies simply use available data and may not appropriately evaluate data quality.

Sources of potential biases in opportunistic occurrence data have also been well-documented in previous work and generally include variation in collection effort and taxonomic, spatial, and temporal biases (4,38–43). Some examples of variables contributing to bias include socioeconomic factors (42,43), the exclusion of common species over rare and flashy ones (44–46), the selection of large and attractive specimens (47), seasonal bias (48), problematic distinction between living and dead-collected specimens and associated post-mortem transportation (49,50), and discarding worn specimens, which results in phenological bias or elimination of specimens with signs of disease (8). Traditional methods for dealing with these issues may include subsampling, data aggregation, and additional surveys (7). Effects of bias can be reduced for certain studies with higher numbers of records, by combining information from different institutions, and including observation records to supplement specimen data (8). Newer statistical and modeling approaches to deal with biases in biodiversity data have also been developed (41,46,51,52). However, it is unclear how often studies actually address issues of error and bias when using opportunistic records.

While several previous studies have reviewed uses of natural history collections data (4,6,8,53), and one study has analyzed field-specific usage for the GBIF index (54), to our knowledge no other study has quantitatively reviewed trends in how species occurrence databases are utilized in published research. Our overarching goal in this study is to determine how such usage has developed since 2010, during a time of unprecedented growth of online data resources. We also determine uses with the highest number of citations, how online occurrence data are linked to other data types, and if/how data quality is addressed. Specifically, we address the following questions:

1. What primary biodiversity databases have been cited in published research, and which databases have been cited most often?
2. Is the biodiversity research community citing databases appropriately, and are the cited databases currently accessible online?
3. What are the most common uses, general taxa addressed, and data linkages, and how have they changed over time?
4. What uses have the highest impact, as measured through the mean number of citations per year?
5. Are certain uses applied more often for plants/invertebrates/vertebrates?
6. Are links to specific data types associated more often with particular uses?
7. How often are major data quality issues addressed?
8. What data quality issues tend to be addressed for the top uses?

## II. LITERATURE SEARCH AND CHARACTERIZATION

We searched for papers that use online and openly accessible primary occurrence records or add data to an online database. Google Scholar (GS) provides full-text indexing, which was important for identifying data sources that often appear buried in the methods section of a paper. Our search was therefore restricted to GS and to the time period of 2010 through the date of the search (April 2017; note when looking at trends over time we remove 2017, as the year was not complete in our dataset). All authors discussed and agreed upon representative search terms, which were relatively broad to capture a variety of databases hosting primary occurrence records. The terms included: *“species occurrence” database* (8,800 results), *“natural history collection” database* (634 results), *herbarium database* (16,500 results), *“biodiversity database”* (3,350 results), *“primary biodiversity data” database* (483 results), *“museum collection” database* (4,480 results), *“digital accessible information” database* (10 results), and *“digital accessible knowledge” database* (52 results)--note that quotations are used as part of the search terms where specific phrases are needed in whole. We downloaded the first 500 records (or all if there were fewer than 500 results), which are presumably the most relevant search returns, for each search term into a Zotero reference management database (55). We obtained citation numbers for each paper from the GS search results at the time of downloading records (April 2017;,56). After removing duplicates across search terms, the final database included 2,500 papers. We then randomly sorted papers into four separate sets of 500 to allow subsampling of the dataset.

For a study to be relevant in this assessment, there must be an indication that the database used is publicly accessible online in a searchable database of biodiversity records. The databases used may include specimen and/or observation-based records from biodiversity data aggregators, online natural history collection databases, websites devoted to capturing citizen science observation records, or newly compiled data that are made available in online databases. Studies were not relevant if they *exclusively* used data that are not available online or from systematic surveys, government monitoring programs, or field data collected explicitly for the study in question. However, papers are relevant if they use these other types of occurrence data *in addition to* online databases of primary occurrence records (see section on data linkages, below), or if they compile these types of occurrence records and deposit them into an existing online biodiversity data aggregator (e.g. GBIF). Twenty-six percent (*n* = 501; see Supplemental File 1 for citation information) of the papers in the final evaluated dataset (*n* = 1,934) were relevant according to these criteria. The full dataset is published and openly accessible (56).

Three of the authors with specialized knowledge of the field (J. Damerow, L. Brenskelle, and R. Guralnick) characterized relevant papers for the first 1000 papers using a standardized tagging protocol based on 14 key topics of interest with over 100 total tags. We developed a list of potential tags and descriptions for each topic; a full list with descriptions of tags is provided in Supplemental Table 1. J. Damerow subsequently checked each tagged paper from the first 1,000 papers to maintain consistency and became the sole tagger for an additional 934 papers. This process allowed the development of a more standardized tagging protocol. The database of tagged papers was then downloaded from Zotero for further data checking and analysis. We used OpenRefine, an open source tool for data cleaning that aggregates similar records for efficient clean-up, to standardize tags from the final dataset.

## III. TRENDS IN USES OF PRIMARY BIODIVERSITY DATA

We characterize a variety of ways in which researchers are using species occurrence records by assessing the prevalence of individual tags corresponding to topics of interest. We identify the most commonly cited databases and most-studied taxa, number of taxa addressed, most common research uses, the types of data most often linked to species occurrence records, and aspects of data quality addressed in these papers. In addition, we determine prevalence of these tags over time to assess positive or negative trends.

### a. Primary biodiversity databases and accessibility of data

We identify 347 primary biodiversity databases used in papers from our dataset (Supplemental Table 2), the URL for each database, and the scale (institution, regional, global, taxa) and regional or taxonomic focus (e.g. Australia, fish) of each database. We then evaluate citation information provided in each paper, and assess whether the data are currently available online or not by visiting associated URLs. The most cited databases include: the Global Biodiversity Information Facility (GBIF), Barcode of Life Data System (BOLDSystem), SpeciesLink, Ocean Biogeographic Information System (OBIS), Australia’s Virtual Herbarium, Tropicos, FishBase, Fishes of Texas, and CONABIO (Table 1).

**Table 1.** Top ten most used biodiversity databases (see Supplemental Table 2 for a comprehensive list).

| Database Name | Number of Papers Citing |
| --- | --- |
| GBIF | 155 |
| BOLDSystems | 27 |
| SpeciesLink | 21 |
| OBIS | 20 |
| Australia's Virtual Herbarium | 19 |
| Tropicos | 16 |
| FishBase | 14 |
| Fishes of Texas | 13 |
| CONABIO | 11 |

Our dataset includes 165 papers that involve compiling and publishing data online (117 data papers and 60 papers that describe a new database, some of these papers overlap). Previous work has outlined best practices for publication of biodiversity data (57–62) and scientific data more generally (e.g. 63). However data are published, primary biodiversity data should also be integrated into an aggregate system with similar data, such as GBIF, OBIS, VertNet, iDigBio, or BoldSystems (62).

Many researchers do not sufficiently cite databases used (64,65), and links to many databases become invalid over time (66–68). We found that 34 percent of papers (*n* =170) had insufficient citation information for one or more databases; this meant that there was either no URL provided to access the database, or the URL was broken. Twenty-six percent of databases (*n* =90) cited in one or more papers from our dataset were totally inaccessible at the time of this assessment. In some cases, researchers appropriately cited a database that is no longer in operation or has subsequently been integrated into an aggregate system. As a result of insufficient data citation practices and lack of data preservation, data are either completely lost or it is impossible to reproduce the dataset used and results. Study reproducibility, strongly linked to data persistence (66), is a key principle in the scientific process and a growing concern across scientific disciplines (e.g. 69). Researchers who have compiled data from multiple sources for a particular analysis can better ensure that their data are accessible and get credit for the work involved in integrating datasets by formally publishing data with descriptive metadata and obtain a persistent DOI (63). The prevalence of inaccessible databases and incomplete database citations indicates that many biodiversity researchers lack the resources to manage and preserve data for the long term and/or are unaware of best practices.

Guidance and infrastructure for citing online data sources have fairly recently emerged and are still evolving (64,70). One major problem is that many papers using biodiversity data have obtained data from an aggregator, such as GBIF, which has potentially drawn from thousands of original data sources. Up to this point, researchers have most often cited GBIF in this case (usually in-text, not in the reference section) and neglect to credit original data sources (65). Even for those who attempt to cite sources, many journals do not allow large numbers of citations in the reference section, and the only solution is to cite sources in a supplement or appendix which does not provide citation credit (65). Data contributors who have submitted data to aggregators are not getting credit for the significant work spent on data management, standardization, and quality control. Ideally, data citations should include DOIs for datasets if they exist and citations of online databases both in text and in the reference section (64,65,71). We will address data citation practices more thoroughly in a separate paper.

### b. Research uses

A primary topic of interest for this work was to characterize research uses of the study databases. An initial list of use tags was developed based on usage outlined in (23), which surveyed needs of primary biodiversity data users. We subsequently split up certain aggregated topics and revised and added use categories based on important subject areas that arose during the tagging process. We ended with 31 potential research use tags, as listed and described in Supplemental Table 1. Most papers had multiple use tags assigned (mean=2.5, max=7). We then determined the average number of citations for papers involving each data use. Number of citations was extracted from the original web snapshots of the Google Scholar searches for each term in April 2017 (56).

Expected trends for research uses in published work include the following: *H1)* Data uses requiring large numbers of dispersed records, such as species distribution models and biodiversity studies, are the most common uses of online databases and have increased over time; *H2)* Data papers and papers describing a new database are likely to have increased in recent years as new venues have grown supporting such publications; and *H3)* Uses involving other online data types (i.e. barcoding, citizen science, species interactions) that can be linked to species occurrence records are likely to increase.

The top research uses for online species occurrence databases—from our dataset of 501 relevant papers—were studies on species distribution (*n*=175), diversity/population studies that usually assess species richness (*n*=122), dataset description (i.e. data papers, *n*=117), taxonomy (*n*=95), conservation (*n*=68), data quality (*n*=68), invasive species (*n*=61), and that described a new database (*n*=60, Fig. 1); see Supplemental Table 1 for full descriptions of each category of research use. The prevalence of most uses did not change from 2010-2016, with the exception of data papers and taxonomy-related studies, which both increased (Fig. 2); taxonomy studies usually involved developing regional species checklists. In the aforementioned survey assessment of user needs for primary biodiversity data (22,23), these same categories of use were among the top ways in which people listed that they use primary biodiversity data. Some exceptions were that a relatively large number of survey respondents claimed that they use data for ecology/evolution studies, natural resources management, life history/phenology studies, and education/outreach, but relatively few published studies used occurrence data for these purposes in our dataset. It is possible that people use data for these purposes, but do not necessarily publish papers on the topic or may not cite databases for this work (72).

**Figure 1.**
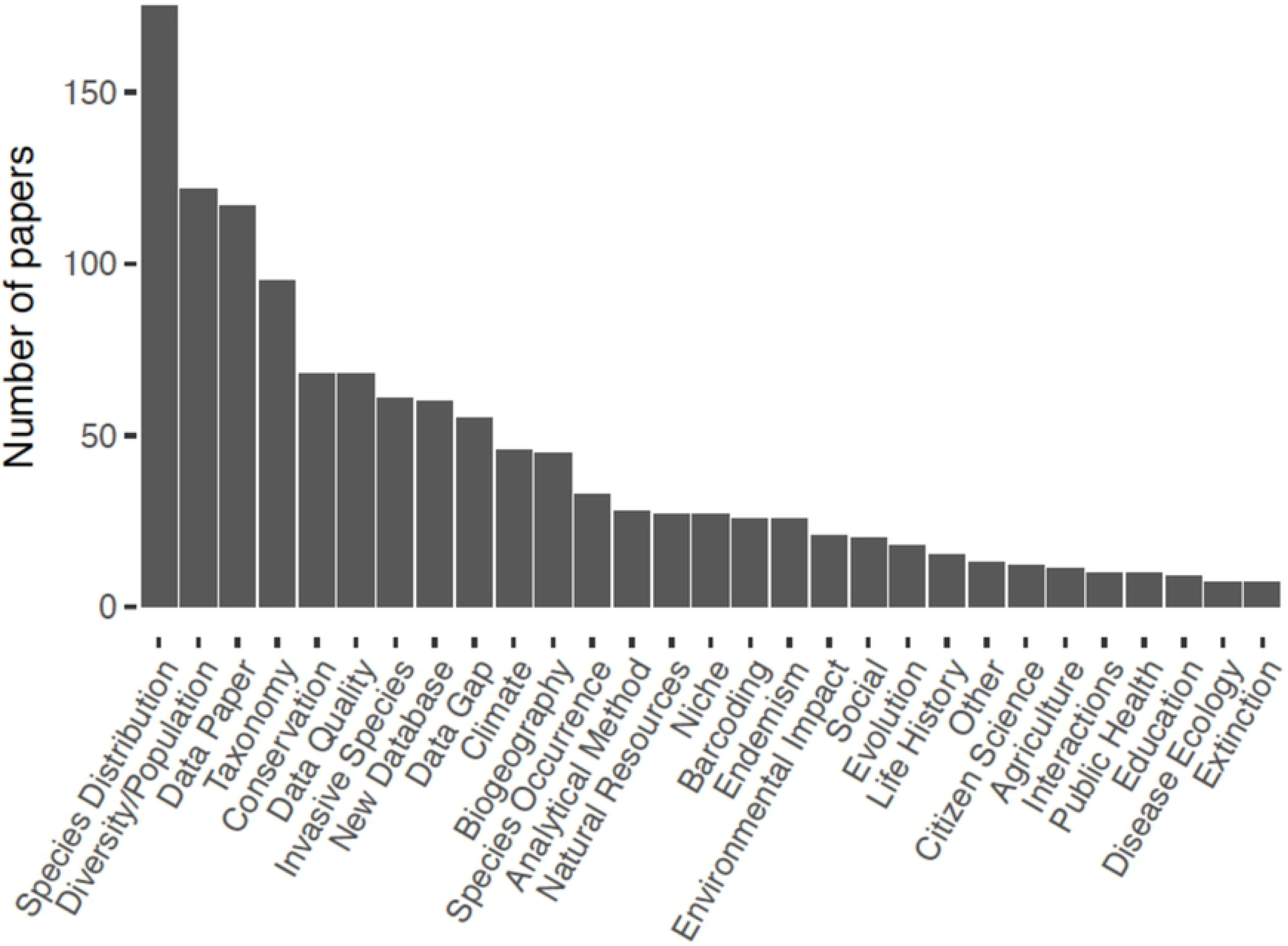
Frequency of major research uses in published papers (*n* = 501) that obtain data from species occurrence records available in online databases. See Supplemental Table 1 for detailed descriptions of each research type.

**Figure 2.**
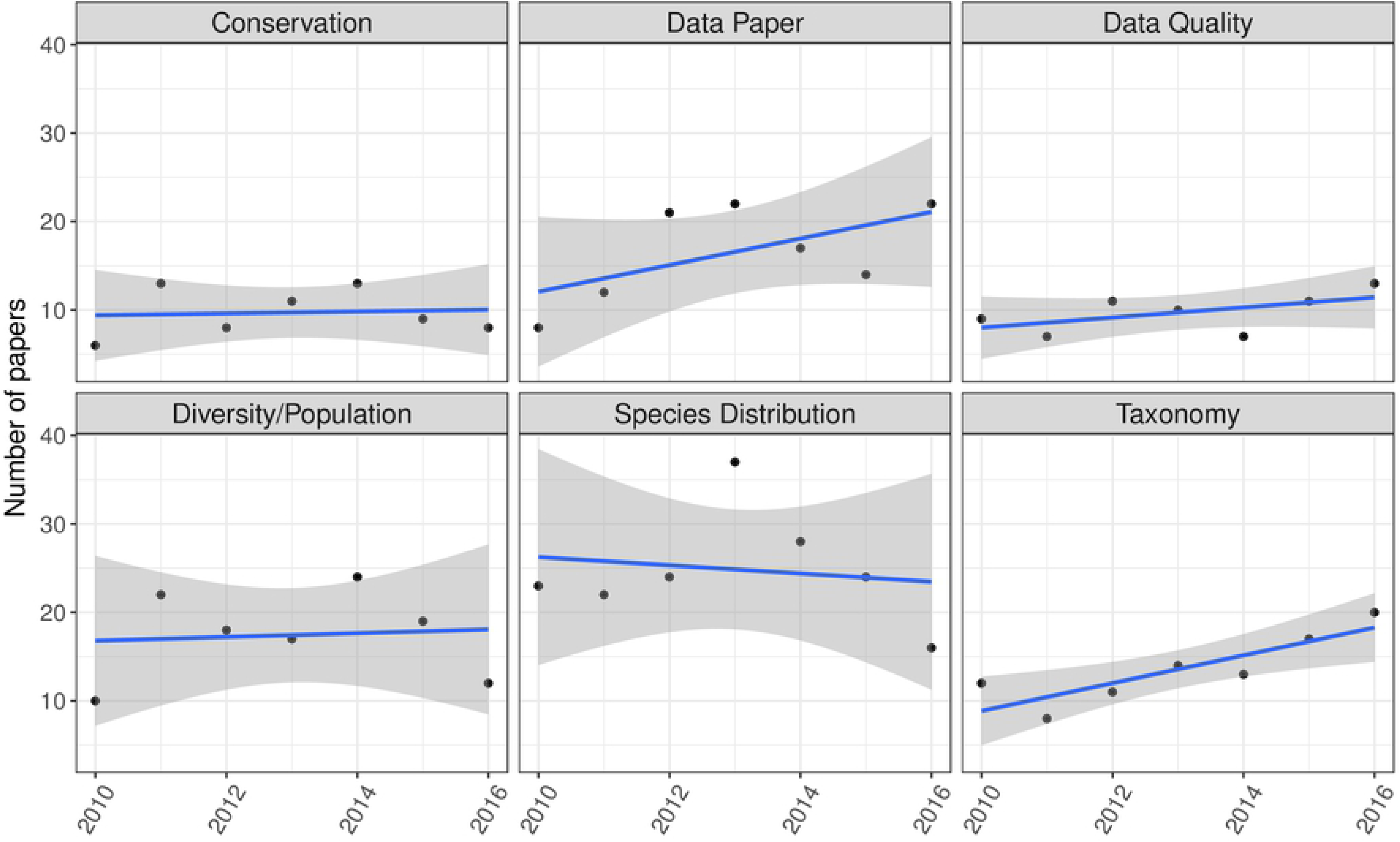
Change in the number of papers from 2010-2016 involving the top six research applications for online species occurrence databases.

Some of the top research uses involved compiling and processing data, as reflected in the high numbers of data papers, papers describing new databases, and papers addressing data quality and data gaps (all of which were among the top ten uses, Fig. 1). The biodiversity community is still in an active stage of compiling existing biodiversity data and dealing with issues of data quality. Data papers and papers describing a new database have increased over time (Fig. 2), which is likely to be the result of the introduction and expansion of many data journals (57,73), online platforms for reporting species occurrence observations such as iNaturalist (74) and eBird (3,75), and efforts over the past decade to digitize specimen records (1,13). More journals accept papers or even focus on publishing high-quality data and recognize this as an important part of the scientific process (62,72,76,77).

Papers with the highest mean number of citations per year involved more applied studies in disease ecology (mean = 18, SD = 33), public health (mean = 8, SD = 7), documenting extinctions (mean = 7, SD = 7), developing a new analytical method to deal with species occurrence data (mean = 7, SD = 8), and citizen science (mean = 7, SD = 6; Table 2). Papers with the highest maximum number of citations per year focused on disease ecology, species diversity, and publishing data (each with a maximum of 97 citations/year; Table 2); we did not account for self-citation here.

**Table 2.** Summary statistics for the number of citations per year for each use of primary biodiversity data. Note that not all papers had citation data available.

| **Data Use** | **n** | **mean** | **sd** | **min** | **max** |
| --- | --- | --- | --- | --- | --- |
| Disease Ecology | 8 | 18 | 33 | 2 | 97 |
| Public Health | 9 | 8 | 7 | 0 | 22 |
| Extinction | 6 | 8 | 7 | 1 | 17 |
| Analytical Method | 26 | 7 | 8 | 1 | 34 |
| Citizen Science | 7 | 7 | 6 | 1 | 17 |
| Species Distribution | 152 | 6 | 10 | 0 | 97 |
| Climate | 46 | 6 | 6 | 0 | 32 |
| Niche | 24 | 6 | 5 | 0 | 20 |
| Data Quality | 59 | 6 | 8 | 0 | 37 |
| Diversity/Population | 108 | 5 | 10 | 0 | 97 |
| Data Paper | 94 | 5 | 11 | 0 | 97 |
| Other(Paleontological) | 3 | 5 | 5 | 0 | 10 |
| Other(Behavior) | 1 | 5 | NA | 5 | 5 |
| Data Gap | 56 | 5 | 6 | 0 | 28 |
| Agriculture | 10 | 5 | 4 | 1 | 13 |
| Invasive Species | 55 | 5 | 5 | 0 | 32 |
| Conservation | 61 | 5 | 6 | 0 | 22 |
| Endemism | 23 | 5 | 5 | 0 | 20 |
| Evolution | 17 | 5 | 3 | 0 | 12 |
| Barcoding | 22 | 5 | 4 | 0 | 16 |
| Biogeography | 41 | 5 | 4 | 0 | 16 |
| New Database | 50 | 4 | 6 | 0 | 29 |
| Species Occurrence | 26 | 4 | 4 | 0 | 22 |
| Interactions | 7 | 3 | 3 | 1 | 9 |
| Natural Resources | 24 | 3 | 3 | 0 | 12 |
| Environmental Impact | 18 | 3 | 2 | 0 | 7 |
| Other(Movement) | 3 | 3 | 2 | 2 | 5 |
| Life History | 10 | 3 | 2 | 1 | 8 |
| Taxonomy | 72 | 2 | 3 | 0 | 16 |
| Other(Ethnobotany) | 1 | 2 | NA | 2 | 2 |
| Education | 5 | 2 | 2 | 0 | 5 |
| Social | 14 | 2 | 1 | 0 | 5 |
| Other(Reference) | 1 | 1 | NA | 1 | 1 |

### c. Taxa addressed

The third major topic for this work was to determine how often different taxonomic groups are represented in papers utilizing biodiversity databases. Taxa in relevant papers were coarsely characterized as plants, vertebrates, invertebrates, fungi, paleo, and/or all taxa; note that we addressed only macro-organisms because they are the focus of non-sequence-based species occurrence databases. These general taxonomic categories also correspond to common divisions for the organization of natural history collections and associated databases. Many papers include more than one taxon, and we use an “all taxa” categorization for studies that use all available data within the species occurrence database(s), such as GBIF. We further categorized taxa addressed in each paper by adding one or more tag(s) for more specific taxonomic classifications (e.g. butterflies, *Danaus plexippus*). While an in-depth assessment of specific taxa is beyond the scope of the current paper, we did tag the number of taxa addressed in each paper, if that number was apparent. Our goals here were to characterize the most commonly studied taxonomic groups, the number of taxa addressed, and to determine uses associated with the three most common organismal groupings (plants, vertebrates, and invertebrates).

Expected trends for taxonomic groups addressed in published work include the following: *H1)* Papers involving plants will be the most common, given work by Tydecks *et al.* (2018); *H2)* Vertebrate data are generally more often applied towards species distribution and conservation studies; *H3)* Invertebrate studies are the least common of the three major groups and are more likely to be the subject of taxonomy, species richness, and barcoding studies; and *H4)* The number of species addressed is likely to increase over time as data for more species become available online and more ambitious projects are undertaken leveraging broad-scale data.

The most commonly studied taxa were plants (*n*=232 papers, 46%), followed by invertebrates (*n*=125, 25%), vertebrates (*n*=124, 25%), “all taxa” (*n*=40, 8%), fungi (*n*=16, 3%), and paleontological specimens (*n*=14, 3%; Table 3). However, the gap between number of papers addressing plants, vertebrates, and invertebrates closed in recent years (2014–2016, Fig. 3). The overall prevalence of plants in this work corroborated a recent bibliometric study, which found that 56% of biodiversity-related papers addressed plants, compared to 29% for vertebrates and 23% for invertebrates (78). The prevalence of plants in the field of biodiversity research may be the result of several factors. Plants are far more diverse than vertebrates (known to be relatively well-studied) and therefore generally require more taxonomic work. Herbarium sheets have also been the easiest historically to digitize, as sheets can be scanned and imaged using more automated processes (11,15). The current prevalence of plants may also partially be the result of a strong history of plant research in Europe; this tendency is known as the “Matthew principle” whereby research concentrates on already well-studied subjects (78). The total number of invertebrate studies was equivalent to the total number of vertebrate studies (Fig. 3). However, invertebrates are much more diverse in terms of species (estimated at 6,755,830 species, see 79), and vertebrates are unquestionably more studied on a per-species basis. The numbers of papers addressing vertebrates and invertebrates has increased slightly and were roughly equivalent over time (Fig. 3). The frequency of papers addressing “all taxa” from online databases has not changed significantly over time (Fig. 3).

**Figure 3.**
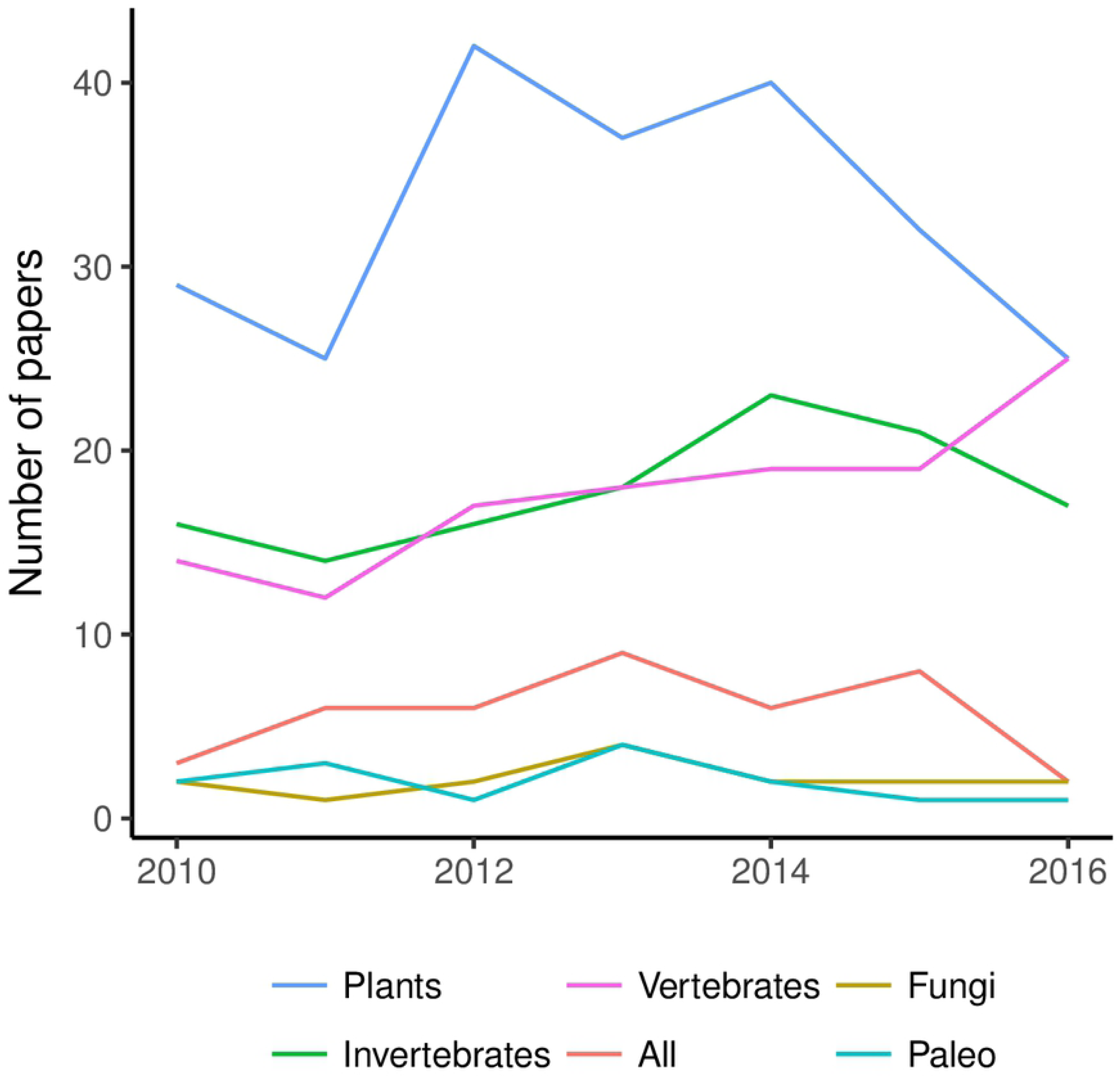
Number of papers addressing the major taxonomic groups and paleontological records.

**Table 3.** Total number of papers from dataset (501) addressing the major taxonomic groups and paleontological specimens.

| Taxa | Number of papers |
| --- | --- |
| Plants | 232 |
| Invertebrates | 125 |
| Vertebrates | 124 |
| All | 40 |
| Fungi | 16 |
| Paleo | 14 |

The most common data uses associated with the major taxonomic groups reflect the general maturity of data products associated with the respective group. Over 50% of vertebrate studies involved investigating species distribution (Fig. 5); vertebrate data are generally more suitable for distribution studies because vertebrates are less diverse, many collections are completely digitized, and data for individual species are likely to contain sufficient numbers of records. Birds in particular have relatively good data available, in part because of online citizen science efforts and associated open data platforms such as eBird (3). While distribution studies were still the most common application for plants and invertebrates, only 33% and 41%, respectively, of plant and invertebrate studies dealt with species distribution. Plants and especially invertebrates are much more diverse, and the average species in these groups are less likely to have data of sufficient quantity and quality to estimate species distribution, although growth in resources especially for plants is closing the gap. Data on insect distributions, in particular, are less complete (or non-existent) for most species and hence may not be suitable for distribution and conservation studies (80,81).

A higher percentage of data papers, taxonomy, and barcoding papers involved invertebrates (Fig. 5), reflecting in part the high taxonomic diversity for this group and need for more data. There are around 60,000 species of vertebrates, an estimated 400,000 plants, and an estimated 5–6 million species of insects—about one million insect species are currently described, which highlights the need for more taxonomic work in this group (19,82). Other invertebrate phyla, such as Mollusca, are highly diverse as well (estimated 70,000–76,000 living species;,83). Digitizing efforts for invertebrates have been particularly challenging, because many clades are so diverse, collections have much larger numbers of specimens, and the typically small specimens are difficult to digitize (84). Automating digitization of such specimens, especially pinned insects and fluid-preserved invertebrates, faces significant obstacles (12,17,85–88).

The use of species occurrence data for conservation followed predicted trends. Vertebrate studies were more likely to address conservation; 23% of papers using vertebrate biodiversity records involved conservation, as compared to 14% of papers using plant records and 12% of papers using invertebrate records (Fig. 5). Twenty percent of vertebrate species are currently classified as threatened, and that number is increasing (89). While vertebrates have more data, they are by no means complete (90); less-studied vertebrates (i.e. fish) are also the least digitized, as compared to birds (91). Large species tend to receive more research focus and conservation funding, and very few conservation assessments exist for invertebrate taxa; most insect species are classified as “data deficient” (e.g. 92). There is much need and potential for using primary biodiversity data to help determine conservation status of insects—perhaps starting with taxa known to be biological indicators of ecosystem health (e.g. 93,94) and insects that provide important ecosystem services (e.g. 95). However, identifying decline requires large numbers of records along with systematically collected surveys over time, which often do not exist for rare and potentially threatened species (96). Opportunistic species occurrence records may therefore be best used to identify data gaps and promising areas for resurveys or standardized long-term monitoring studies when dealing with species decline (46).

Contrary to expectations, we found that studies addressing “all taxa” remained fairly consistent over time (Fig. 3), and the maximum number of taxa addressed did not increase (Fig. 4). However, this may simply be an effect of small sample sizes. Only four papers involved numbers of species in the hundreds of thousands over the period of 2010-2017 (Table 4). Most papers focused on numbers of species in the single or double digits (Table 4). We found that the top data uses for papers that addressed “all taxa” involved data compilation and data quality (data quality assessments, data gap studies, data papers, and reporting on new databases, respectively). We argue that the scale of data that needs processing, along with issues of often sparse data, data obsolescence (97), and data of uncertain quality, make large-scale analyses challenging for anyone but a small group of data sciences-savvy end users. Additionally, effective large-scale assessments are often impossible without significant investments and active collaboration across study domains (e.g. taxonomy, ecology, biodiversity informatics) and geographical regions (98).

**Figure 4.**
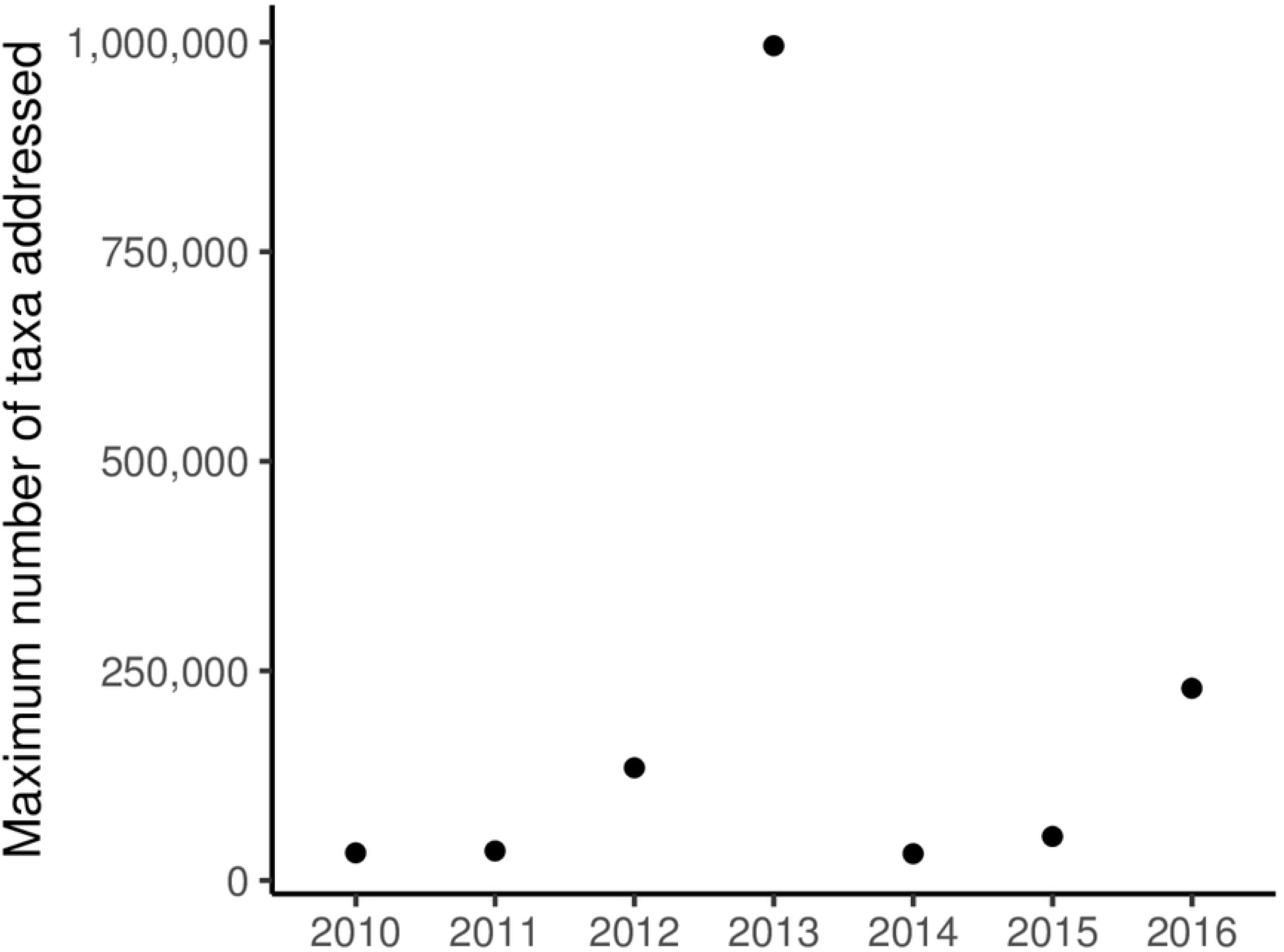
Maximum number of taxa addressed in papers (*n*=501) from 2010-2016.

**Figure 5.**
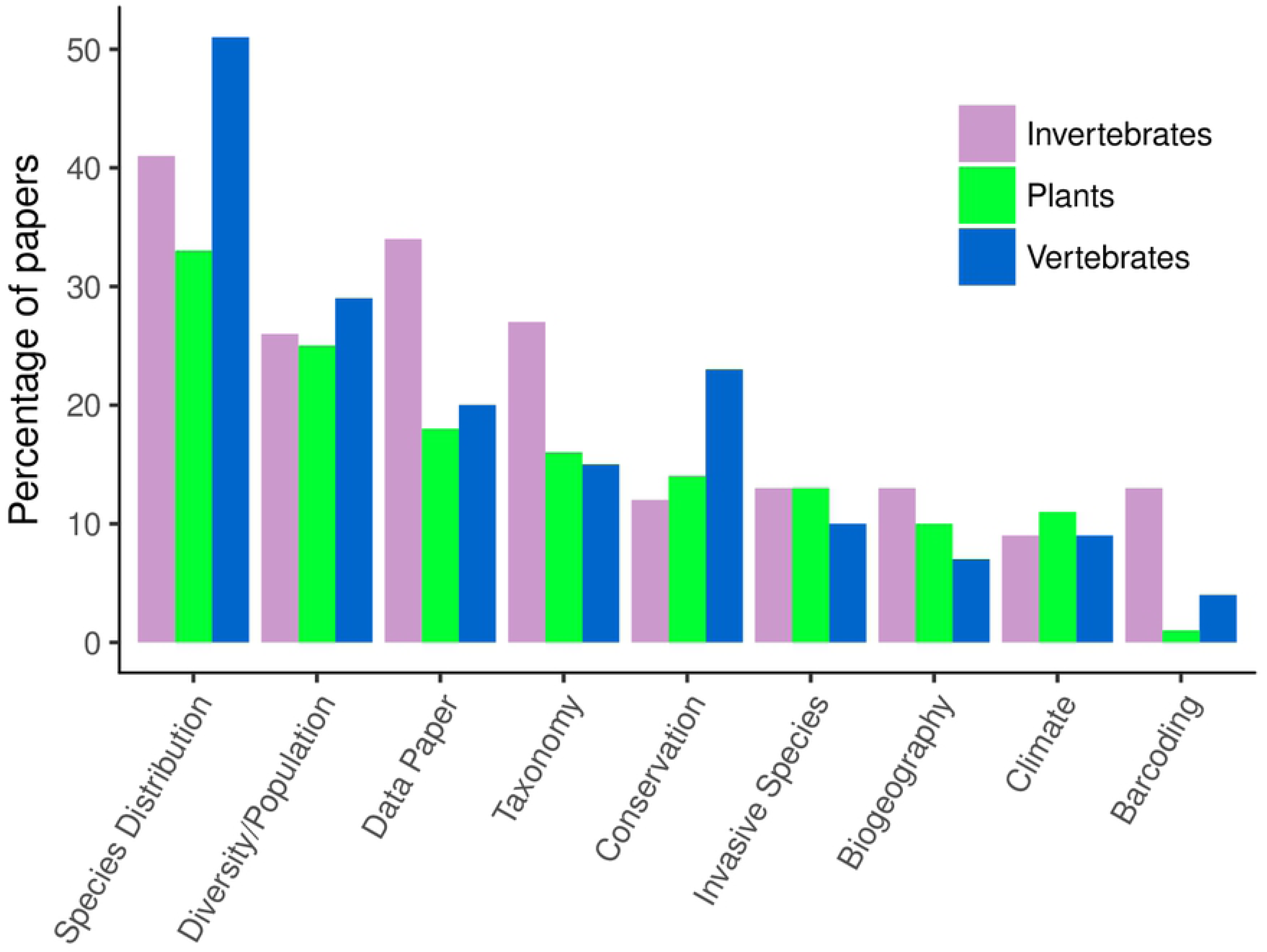
Percentage of papers involving each of the major taxonomic groups (invertebrates, plants, and vertebrates) that use species occurrence databases for certain research applications: species distribution, diversity/population, data paper, taxonomy, invasive species, biogeography, climate change, and barcoding.

**Table 4.** Number of taxa addressed by papers using online species occurrence records.

| Number of taxa addressed | Number of papers |
| --- | --- |
| 1-9 | 113 |
| 10-99 | 106 |
| 100-999 | 82 |
| 1,000-9,999 | 68 |
| 10,000-99,999 | 22 |
| 100,000-999,999 | 4 |

### d. Links to other data types

We determine how studies link primary biodiversity data to other data types by characterizing the variety of data compiled and used in each study (see Supplemental Table 1 for full descriptions of 28 data linkage tags). We searched for information regarding other data types used within the methods section of each paper. Data link tags fall under four general categories of data types, including 1.) other types of occurrence data (i.e. data from literature, field surveys, species catalogues, private data); 2.) attributes of species occurrence data (e.g. information about the holding collections of specimens, species traits, conservation status, genetic data, associated image(s), species interactions, population data); 3.) environmental data (e.g. climate, geographic information, habitat, ecoregion, etc.); and 4.) data that can be used to determine biases or gaps (socioeconomic data, expert knowledge, and accessibility of sites—with the last usually evaluated through proximity to roads or research institutions). We then determine the average number of data link tags associated with the six top uses, and the most common data type associated with each of these top uses.

Expected trends for studies using other data types linked to species occurrence records: *H1*) Climate data are likely to be the most common environmental variable linked to species occurrence records; and *H2*) Other types of occurrence data are also commonly used, as studies often need more data records than are currently available.

Data types that were most often used in association with online species occurrence databases (out of 501 relevant papers) included occurrence records from previously published literature (*n*=189), climate (*n*=149), occurrence records from surveys (*n*=143), collection information (*n*=135), habitat (*n*=118), traits (*n*=111), and geographic data (*n*=106, Fig. 6). Three data types increased from 2010–2016, including collection, genetic, and phylogenetic data (Fig. 6). The average number of data linkages per paper was four (ranging from one to 11).

**Figure 6.**
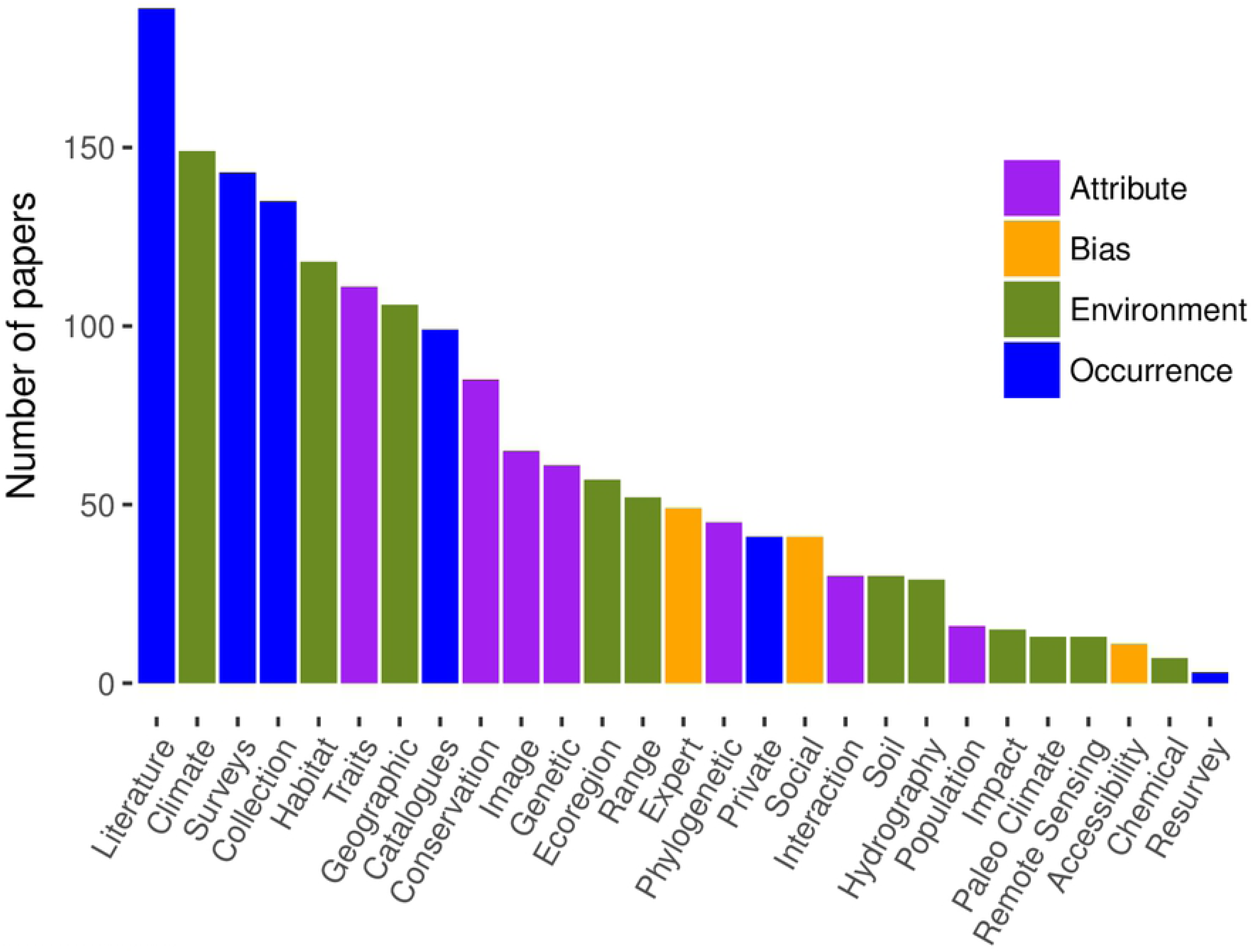
Number of papers that incorporate other data types to supplement or associate with online species occurrence records. Data types fall within one of four categories, including 1.) attributes of occurrence information, 2.) data types that may help address bias in the data, 3.) environmental variables, and 4.) other kinds of occurrence data.

Table 5 summarizes top data linkages for different key uses. As predicted, climate is often a critical data linkage, especially for species distribution where it is the most common linkage, and for diversity/population studies where it is a close second. For data papers and taxonomy studies, both collection data and literature data were often the most common data linkages. Conservation-focused studies that included species occurrences from databases also linked conservation status, habitat, literature, and climatic data. Data quality studies often included a variety of data linkages, with little sorting of top linkages likely representing the high dimensionality of data quality issues.

**Table 5.**
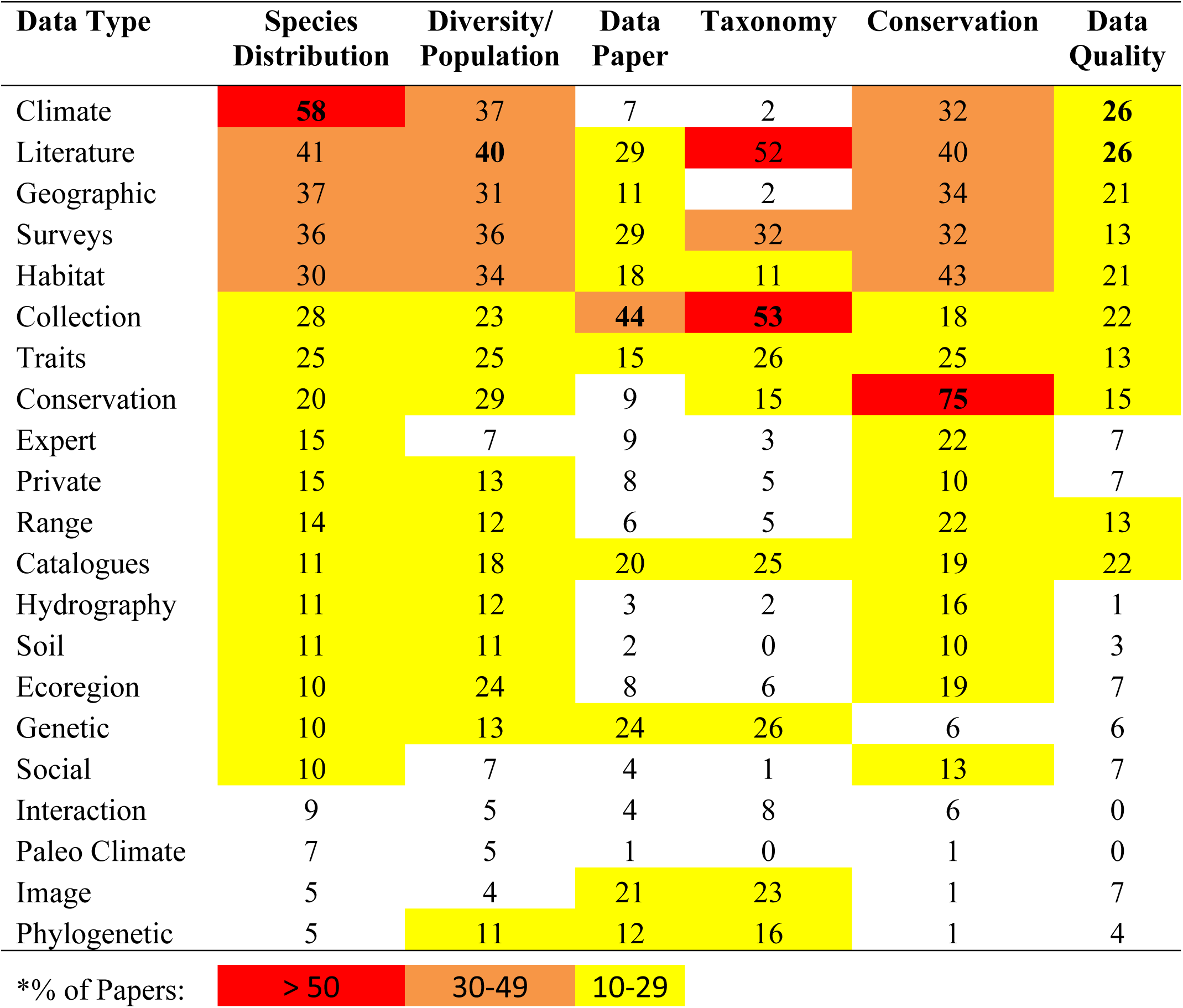
Percentage of papers that associate online occurrence data with other data types— separated by the six top uses of these databases. Nine data types with the lowest percentages were removed from table. The top data type for each research use is bolded, and percentage values above 10% are highlighted yellow, orange, and red*.

The high prevalence of studies compiling occurrence records from other sources indicates a continued demand for more and continued specimen sampling and the need for more progress in getting these data digitally captured and into online databases (i.e. data papers and new database development). Three of the top five data types linked to online occurrence records were other types of occurrence data, including literature-based occurrence data, surveys, and specimen data from natural history collections (*n*=189, *n*=145, and *n*=135 papers used these data types, respectively). Sometimes the compiled data eventually make it into online data aggregators, such as GBIF, and sometimes they do not. Continued advocacy for data publication will be important to maximize the potential use of all biodiversity data.

Environmental data used in conjunction with online biodiversity records are usually applied in studies of species distribution. Specific environmental parameters used to predict distribution should be informed by expert knowledge of the requirements of a given species. Among environmental variables, climate data are perhaps the most readily available, relevant for the distribution of organisms on a global scale, and provide essential information for determining impacts of climate change on distribution (99,100). Our data show that climate is indeed the most common environmental variable used in association with occurrence records (Fig. 6; also documented in 54). The second and third most common environmental data types used were geographic and habitat, which usually included GIS layers for elevation and land use and/or vegetation (see Supplemental Table 1). Elevation, land use, and vegetation data are also among the most readily available environmental data types, and are often relevant for evaluating species distribution at smaller spatial scales (101). Despite increasing calls for incorporating relevant biotic interactions into models, only nine distribution studies incorporated data on interactions (i.e. competitive, consumptive, symbiotic, or pathogenic relationships), and 30 studies overall involved species interactions. The relatively low prevalence of species interaction information in these studies is thought to be primarily due to the large spatial scales usually considered in distribution models. Biotic interactions are often studied on a smaller scale by community ecologists, while distribution modeling is often done by macroecologists (102). Primary species occurrences may provide needed data for studying biotic interactions on a larger scale, but these data are often not digitized, even if they exist in collections, and compiling data of sufficient quantity and quality for a given taxon remains an obstacle due to lack of automated data capture options for invertebrate collections.

The only data types that have increased over time were specimen collection, genetic, and phylogenetic data (Fig. 7). We expected to see an increase in use of genetic data in particular, as these data are known to have expanded with the growth of databases such as the Barcode of Life Data System (BOLDSystems), linking molecular, morphological, and distribution data (103); the number of records in BOLDSystems increased from about 0.5 million in 2007 to 1.5 million today (104). Further, large-scale phylogenetic resources such as Open Tree of Life (105), launched in 2015, have made it easier than ever before to assemble those resources with other species data. The increasingly available collections, genetic, and phylogenetic data are highly relevant in taxonomy-related studies and data papers, which increased over time (Fig. 2).

**Figure 7.**
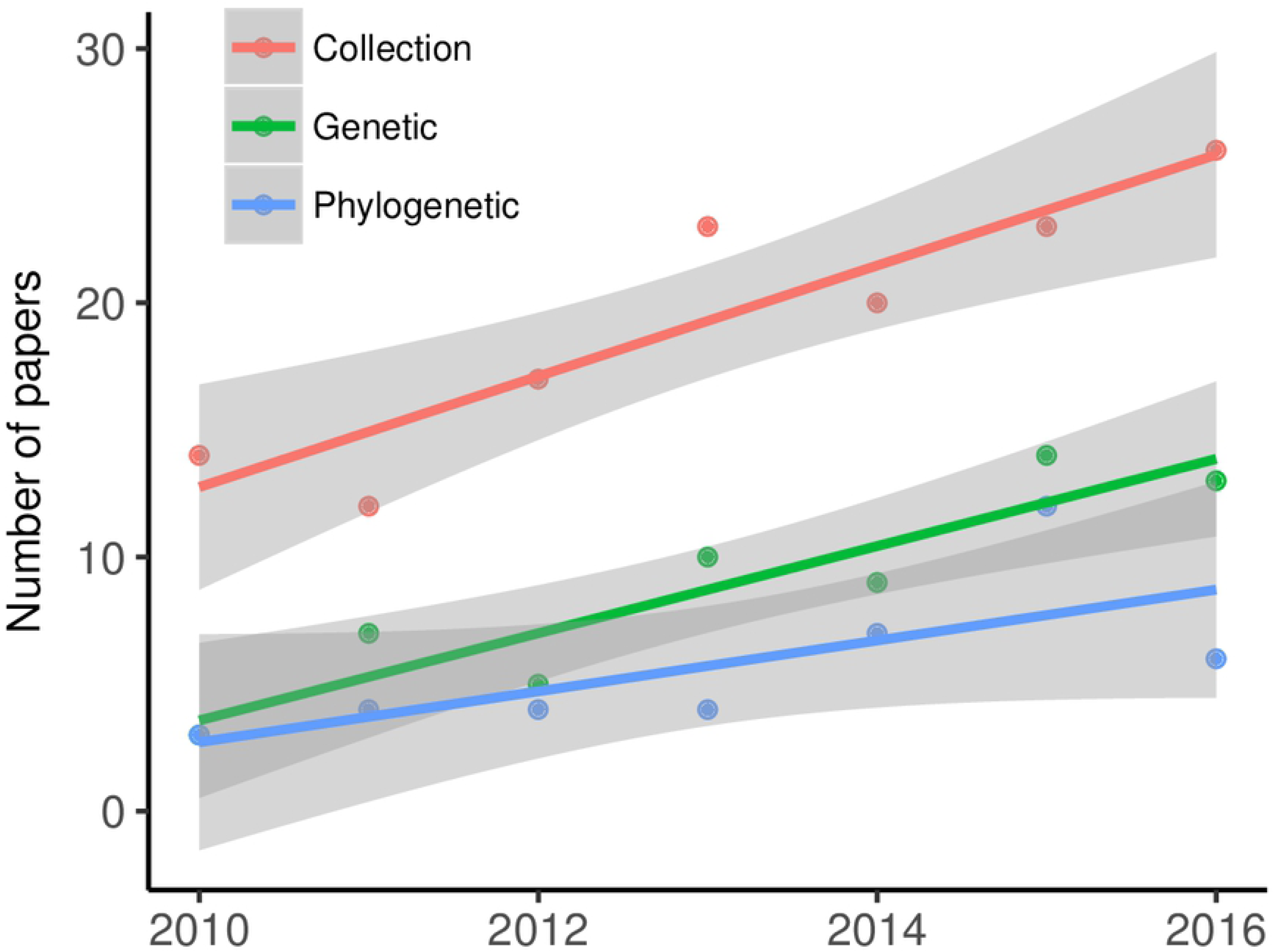
Data types that increased over the period from 2010 through 2016. These include data needed for taxonomic/phylogenetic studies, namely those from natural history specimens, genetic data, and phylogenetic data.

Both taxonomy and data papers used collection data most frequently in addition to data already available in online databases. Taxonomy uses of online species occurrence databases sometimes involve describing new species, but more commonly involve compilation of regional species checklists. The most traditional use of collections data is for taxonomy, so it is not surprising that over 50% of taxonomy papers also involve collections and literature data. The relatively high percentage of data papers that involve collections data (44%) reflects recent digitization efforts for natural history collections (1,9,13,106).

### e. Data quality

We characterize papers that address major data quality issues known to be associated with species occurrence data, including both common errors and biases. Data quality tags involve improving data quality for a particular purpose addressed in the paper. Taxonomic nomenclature, species identification, spatial, and temporal data quality tags represent adjustments to the dataset used in a study that at least partially corrects the associated errors (see Supplemental Table 1). We also characterize studies that exclude certain inappropriate records, remove records with high georeferencing uncertainty, remove outliers, and those that address collection effort—see Supplemental Table 1). In addition to errors, some studies address specific biases known to be a problem in opportunistic datasets, including taxonomic, spatial, temporal, and environmental biases. Finally, we have a “detection” tag to represent use of statistical methods to estimate detection probability (51). We assess the average number of quality tags associated with papers overall, and the most common data quality issues addressed within each of the top uses. We hypothesize that the most common data quality issues addressed are likely to be checks for correct taxonomic nomenclature and correct georeferences.

Overall, 69% of studies from our dataset that used online species occurrence records addressed one or more aspects of data quality. The biggest data quality concerns cited by users of primary biodiversity data in a recent survey (23) were georeference quality and taxonomic quality—we found that studies addressed these issues in 24% (spatial error in georeferences), 39% (taxonomic nomenclature), and 19% (species identifications) of published papers from our dataset (Table 6). Two data quality checks increased from 2010 to 2016: correcting taxonomic nomenclature and specimen identification (Fig. 8), reflecting also the increase in taxonomy-related and data papers.

**Figure 8.**
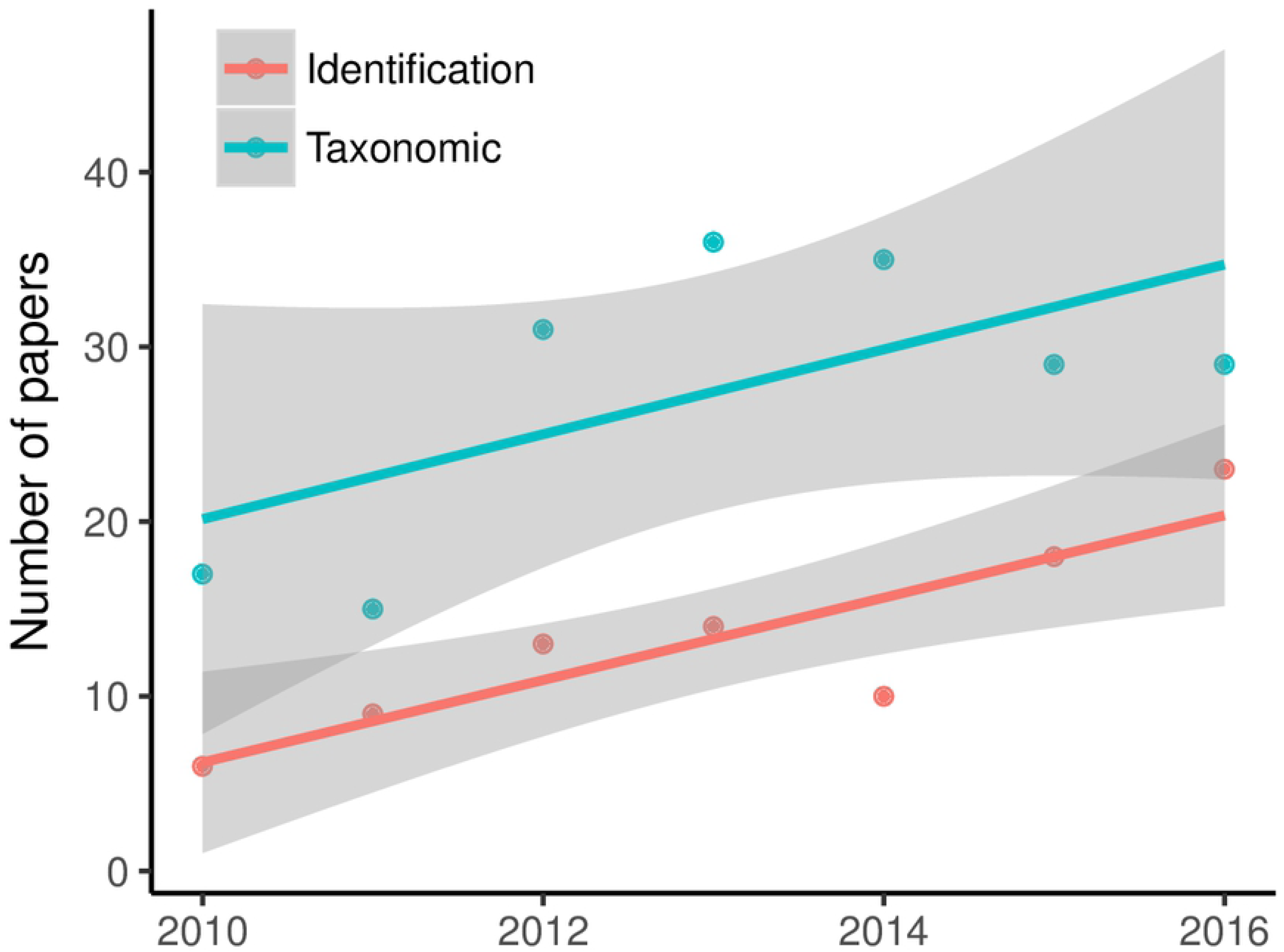
Number of papers that address identification errors and/or update taxonomic nomenclature over the period of 2010-2016.

**Table 6.** Papers from dataset (*n* = 501) that addressed data quality issues associated with species occurrence records.

| Quality Tag | Number of Papers | Percentage |
| --- | --- | --- |
| Taxonomic | 193 | 39% |
| Spatial | 121 | 24% |
| Identification | 94 | 19% |
| Spatial Bias | 59 | 12% |
| Exclusion | 57 | 11% |
| Effort | 50 | 10% |
| Precision | 30 | 6% |
| Temporal | 18 | 4% |
| Outliers | 17 | 3% |
| Temporal Bias | 11 | 2% |
| Taxonomic Bias | 9 | 2% |
| Environmental Bias | 6 | 1% |
| Detection | 4 | 1% |

Spatial errors and taxonomic nomenclature are generally the easiest data quality errors to correct. Non-experts can check for spatial outliers or incorrect georeferences using standardized methods and online georeferencing tools (35,107). Depending on data needs, one may also use existing error radii associated with georeferenced coordinates to select appropriate records for a study. However, most records in GBIF, for example, still do not have error radii; in a recent assessment of GBIF records for Odonata, Ephemeroptera, Plecoptera, and Trichoptera from the U.S.A., we found that the percentage of records with error radii associated with them was only 7-36% for these aquatic insect groups (as of April 2017). Of the 6.2 million catalogued molluscan lots in U.S. and Canadian collections, 4.5 million have undergone some form of data digitization. Of these, about 1.1 million (24%) of digitized records have been georeferenced, which represents 18% of all catalogued lots (47). However, only a subset of these have error radii associated. Many digitization efforts for insects in particular have prioritized transcribing and publishing specimen label information and have not yet begun or completed georeferencing.

Online taxonomic catalogues and tools to check records against updated catalogues are available for correcting taxonomic nomenclature (108,109). However, we still have not reached the major goal of having online taxonomic data sources that are consistently updated by taxonomic experts for all species, although community-supported resources such as FishBase (110), WoRMS (111), and the latter’s affiliated databases such as MilliBase (112), and MolluscaBase (113) are approaching that goal for many taxonomic groups. Other groups may lack online sources or have sources that are significantly out of date (114). Unfortunately, the decline in resources devoted to the field of taxonomy does not bode well for achieving a unified taxonomic backbone usable for resolving all taxonomic issues (115,116). Given the speed of taxonomic concept changes (117), lack of updated resources is a significant impediment to proper data integration. The best way for taxonomic experts to help ensure that nomenclature for their group is current is to engage with the community-supported and specialist-edited taxonomic database projects in their respective fields. The combined data of massive authority file efforts spanning multiple taxon groups, such as those covered by WoRMS, allow for novel approaches to data analysis (118).

Correcting species identifications requires taxonomic expertise for many organisms, particularly high-diversity groups such as insects. Many users outside of the community of trained collection scientists may not understand or be interested in taxonomic concepts (1). Therefore, despite misidentification being a well-known problem, this issue is less often directly addressed in papers. For those who are not taxonomic experts, some possible approaches to address misidentifications include: choosing taxonomic groups that are relatively easy to identify and less likely to have identification error, or including only records identified by reliable experts. For broad-scale biodiversity studies it may be appropriate to check occurrence locations against known ranges (where those exist); one may then identify outliers in the data where species are found in regions where they are not known to occur. Such efforts require both taxonomic and geospatial skills, although some automation may be possible (119).

Biases that result from variation in collection effort across space, time, taxonomic groups, and environments are also well-known problems in opportunistic biodiversity records (30,39,40,80). The most commonly addressed bias in our dataset was spatial (addressed in 12% of papers, Table 7), as it is important for accurate species distribution modeling, and some methods to deal with spatial bias have been developed (39). Other forms of bias were rarely addressed in only 1–2% of papers and include temporal bias (usually seasonal bias for certain times of year, or bias for certain years where specialists are active), taxonomic bias (e.g. preference for endangered species, charismatic taxa, avoiding common species or pests)(45), and environmental bias (e.g. preference for collecting in certain habitats or climates) (39).

**Table 7.**
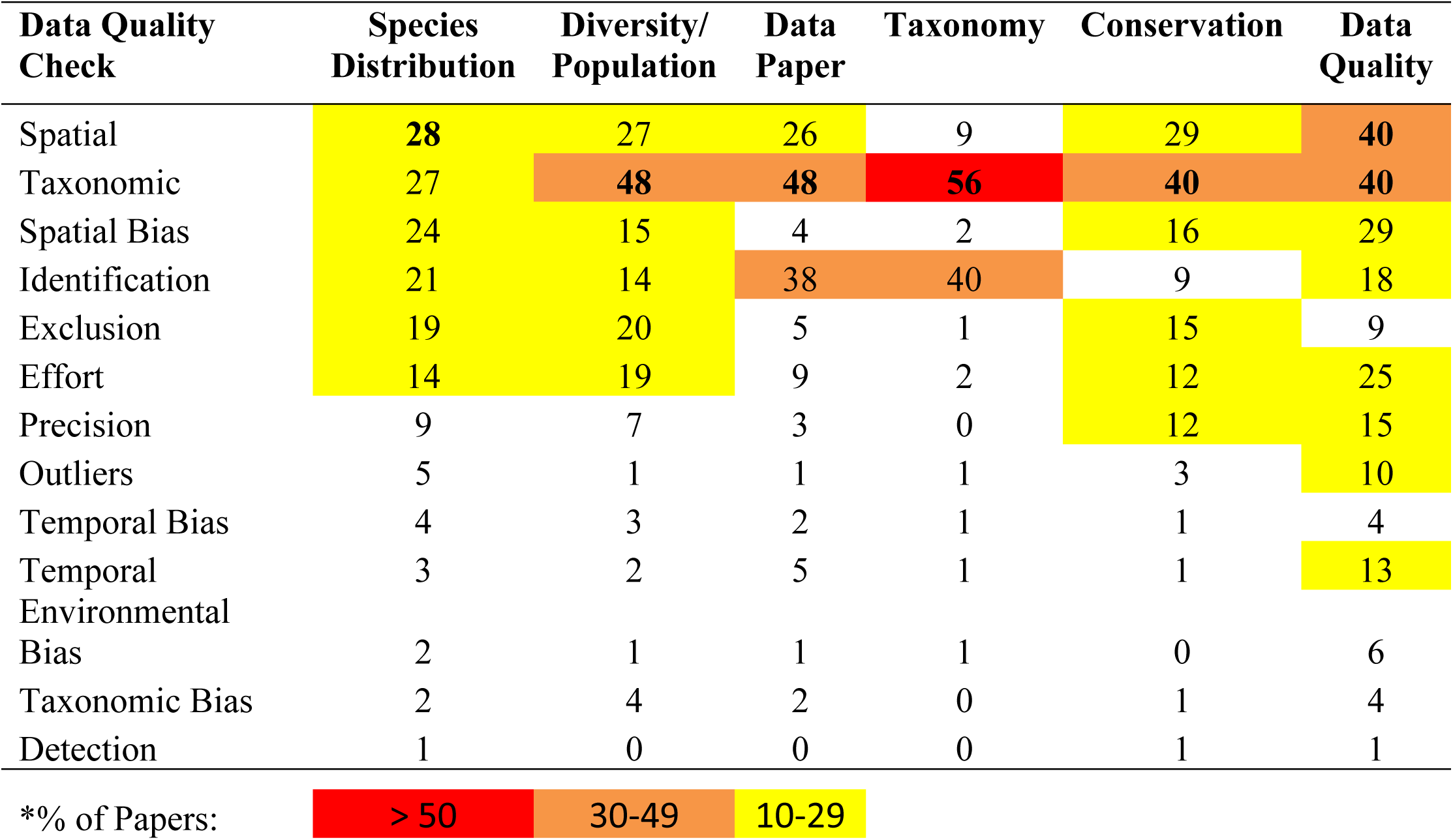
Percentage of papers that check aspects of data quality for online occurrence data—separated by the six top uses of these databases. Nine data types with the lowest percentages were removed from table. The top data type for each research use is bolded, and percentage values above 10% are highlighted yellow, orange, and red*.

Data quality issues addressed are often dictated by the specific use. The most commonly checked data quality issues for papers involving species distribution were spatial errors (28% of distribution studies), taxonomic nomenclature (27%), spatial bias (24%), specimen identification (21%), and excluding inappropriate records (19%; Table 6). Taxonomic nomenclature was the most commonly checked data quality issue for all other top uses, ranging from 40% of papers (conservation and data quality uses) to 56% (taxonomy). In general, taxonomy papers only check issues related to nomenclature and identification. Data quality papers tend to focus evenly on the two most easily corrected issues (spatial and taxonomic, each 40% of data quality papers), followed by accounting for spatial bias (29% of data quality papers), effort (25%), and correcting specimen identification (18%). Diversity/population and conservation papers both also address taxonomic nomenclature and spatial errors most frequently (Table 7).

Automated data quality annotations are growing within the major online data aggregators (e.g. GBIF, iDigBio), but there is still much room to improve upon methods to easily tag data and highlight errors, biases, and uncertainty levels in the data. We need better methods to document confidence in data at a record and dataset level (22). When data quality is addressed, it is usually done manually, and workflows are difficult to document, extend, and share. More recently, programs to automate and document data cleaning workflows have been developed, such as Kurator, a Kepler data curation package (36), but are not yet widely used due to the highly technical user interface, and have uncertain future support. Biodiversity databases allow efficient access to data that can expedite work, but care is still needed when using these resources. Data quality improvements on a large scale will require additional investment in data enhancements (e.g. collaborative georeferencing using standardized point-radius method) and quality control (e.g. efficiently identifying records that may need correction or attention from taxonomic experts).

## IV. CONCLUSIONS AND NEXT STEPS

(1) A high proportion of studies did not sufficiently cite databases, and many databases were no longer accessible at the time of this study; in most cases it was unclear whether the data were lost or moved to an aggregator. Continued efforts in data preservation and promoting best practices in data citation are essential for advancing scientific reproducibility, sustaining data resources, and encouraging publication of high-quality biodiversity data.
(2) The increasing number of data papers over time reflects progress in digitization and online platforms for reporting observations through citizen science, as well as increases in journals that support data publication. Continued growth of data publications will enhance the efficiency and relevance of the field in addressing biodiversity conservation and environmental management.
(3) Our study corroborated a recent bibliometric analysis of the larger field of biodiversity research, finding that more studies address plants (46% of studies using biodiversity databases) than vertebrates (25%) and invertebrates (25%). The prevalence of plants in studies that use online biodiversity databases may be due to a strong history of plant diversity work in Europe in particular, and the relative ease with which herbarium records can be digitized by scanning herbarium sheets.
(4) While studies overall were less common for vertebrates than for plants, vertebrates may generally be more suitable for distribution studies because the group is less diverse, many collections are completely digitized, there are prolific citizen science communities reporting bird observations in particular, and data for individual species are more likely to contain sufficient numbers of records. Conservation studies are also more common for vertebrates, likely because they are disproportionately represented in threat assessments. In contrast, highly diverse invertebrates are more likely to be the subject of foundational biodiversity studies, such as taxonomy, barcoding, and data papers.
(5) It is concerning that a relatively large proportion of studies does not explicitly address data quality—only 69% of studies in our dataset reported addressing one or more aspects of data quality. Authors who do address data quality are most likely to standardize nomenclature using online resources or to correct spatial errors. For nearly all uses of these data, there are errors and biases that can compromise results when using opportunistic records. Improving upon automated solutions to flag errors, and efficient mechanisms to report and correct data quality issues is critical in advancing the relevance and broadest use of this type of biodiversity data (120).
(6) Significant investments in data enhancement and quality control are needed. This may be one limiting factor holding back studies that utilize all data currently held within biodiversity databases and studies that address very large numbers of taxa within clades. We found only four studies since 2010 that address hundreds of thousands of taxa, and most papers address numbers of taxa in the single or double digits. Large-scale improvements in data availability and fitness will require interdisciplinary effort and collaboration.
(7) To limit the scope of the present paper, we focused efforts here on data citation, research uses, general taxa addressed, data linkages, and data quality issues addressed. However, we are also utilizing the dataset of tagged papers to address additional questions regarding author connectedness and collaboration across institutions, countries, and disciplines. Such next-step efforts will provide additional context about the nature and scope of collaborations and resources that coalesce around digitally accessible primary biodiversity data.

## V. ACKNOWLEDGEMENTS

This research was supported in part through a Bass Postdoctoral fellowship to J. Ball-Damerow at the Field Museum of Natural History (Chicago, USA), under the mentorship of P. Sierwald and R. Bieler, and by the Negaunee Foundation. We also thank Paula Zermoglio, and John Wieczorek for their advice and assistance in developing methodology during the initial stage of this work.

## SUPPORTING INFORMATION

**Supplemental Table 1.** Description of tags used to characterize papers, and number of papers assigned to each tag.

**Supplemental Table 2.** Online biodiversity databases cited in published research and information on database scale, accessibility, and subject focus of the database (region, institution, and/or taxa included).

**Supplemental File 1.** File in csv format containing citation information for 501 relevant journal articles analyzed in this review.

## REFERENCES

1. Beaman R, Cellinese N. Mass digitization of scientific collections: New opportunities to transform the use of biological specimens and underwrite biodiversity science. ZooKeys. 2012 Jul 20;209:7–17.

2. Matsunaga A, Thompson A, Figueiredo RJ, Germain-Aubrey CC, Collins M, Beaman RS, et al. A Computational- and Storage-Cloud for Integration of Biodiversity Collections. In: 2013 IEEE 9th International Conference on e-Science. 2013. p. 78–87.

3. Sullivan BL, Aycrigg JL, Barry JH, Bonney RE, Bruns N, Cooper CB, et al. The eBird enterprise: an integrated approach to development and application of citizen science. Biol Conserv. 2014;169:31–40.

4. Shaffer HB, Fisher RN, Davidson C. The role of natural history collections in documenting species declines. Trends Ecol Evol. 1998 Jan 1;13(1):27–30.

5. Ristaino JB. Tracking historic migrations of the Irish potato famine pathogen, Phytophthora infestans. Microbes Infect. 2002 Nov 1;4(13):1369–77.

6. Suarez AV, Tsutsui ND. The Value of Museum Collections for Research and Society. BioScience. 2004 Jan 1;54(1):66–74.

7. Graham CH, Ferrier S, Huettman F, Moritz C, Peterson AT. New developments in museum-based informatics and applications in biodiversity analysis. Trends Ecol Evol. 2004 Sep 1;19(9):497–503.

8. Pyke GH, Ehrlich PR. Biological collections and ecological/environmental research: a review, some observations and a look to the future. Biol Rev. 2010;85(2):247–266.

9. Baird RC. Leveraging the fullest potential of scientific collections through digitisation. Biodivers Inform [Internet]. 2010 Oct 9 [cited 2016 Aug 16];7(2). Available from: https://journals.ku.edu/index.php/jbi/article/view/3987

10. GBIF [Internet]. [cited 2019 Apr 5]. Available from: https://www.gbif.org/

11. Baker B. New Push to Bring US Biological Collections to the World’s Online Community Advances in technology put massive undertaking within reach. BioScience. 2011 Sep 1;61(9):657–62.

12. Blagoderov V, Kitching I, Livermore L, Simonsen T, Smith V. No specimen left behind: industrial scale digitization of natural history collections. ZooKeys. 2012 Jul 20;209:133–46.

13. Page LM, MacFadden BJ, Fortes JA, Soltis PS, Riccardi G. Digitization of Biodiversity Collections Reveals Biggest Data on Biodiversity. BioScience. 2015 Sep 1;65(9):841–2.

14. Ariño A. Putting your Finger upon the Simplest Data. Biodivers Inf Sci Stand. 2018 Jun 15;2:e26300.

15. Nelson G, Paul D, Riccardi G, Mast A. Five task clusters that enable efficient and effective digitization of biological collections. ZooKeys. 2012 Jul 20;209:19–45.

16. Tulig M, Tarnowsky N, Bevans M, Kirchgessner A, Thiers B. Increasing the efficiency of digitization workflows for herbarium specimens. ZooKeys. 2012 Jul 20;209:103–13.

17. Hudson LN, Blagoderov V, Heaton A, Holtzhausen P, Livermore L, Price BW, et al. Inselect: Automating the Digitization of Natural History Collections. PLOS ONE. 2015 Nov 23;10(11):e0143402.

18. Allan EL, Livermore L, Price B, Shchedrina O, Smith V. A Novel Automated Mass Digitisation Workflow for Natural History Microscope Slides. Biodivers Data J. 2019 Jan 3;7:e32342.

19. Pimm SL, Jenkins CN, Abell R, Brooks TM, Gittleman JL, Joppa LN, et al. The biodiversity of species and their rates of extinction, distribution, and protection. Science. 2014 May 30;344(6187):1246752.

20. Alroy J. Current extinction rates of reptiles and amphibians. Proc Natl Acad Sci. 2015;112(42):13003–13008.

21. Régnier C, Achaz G, Lambert A, Cowie RH, Bouchet P, Fontaine B. Mass extinction in poorly known taxa. Proc Natl Acad Sci. 2015;112(25):7761–7766.

22. Faith D, Collen B, Ariño A, Koleff PKP, Guinotte J, Kerr J, et al. Bridging the biodiversity data gaps: Recommendations to meet users’ data needs. Biodivers Inform. 2013;8(2). Available from: https://journals.ku.edu/index.php/jbi/article/view/4126

23. Ariño AH, Chavan V, Faith DP. Assessment of user needs of primary biodiversity data: Analysis, concerns, and challenges. Biodivers Inform [Internet]. 2013 Jul 9 [cited 2016 Nov 14];8(2). Available from: https://journals.ku.edu/index.php/jbi/article/view/4094

24. Guralnick R, Hill A. Biodiversity informatics: automated approaches for documenting global biodiversity patterns and processes. Bioinformatics. 2009 Feb 15;25(4):421–8.

25. Sousa-Baena MS, Garcia LC, Peterson AT. Knowledge behind conservation status decisions: data basis for “Data Deficient” Brazilian plant species. Biol Conserv. 2014;173:80–89.

26. Feeley K. Are We Filling the Data Void? An Assessment of the Amount and Extent of Plant Collection Records and Census Data Available for Tropical South America. PLOS ONE. 2015 Apr 30;10(4):1–17.

27. Meyer C, Kreft H, Guralnick R, Jetz W. Global priorities for an effective information basis of biodiversity distributions. Nat Commun [Internet]. 2015 Dec [cited 2018 May 24];6(1). Available from: http://www.nature.com/articles/ncomms9221

28. Beck J, Ballesteros-Mejia L, Buchmann CM, Dengler J, Fritz SA, Gruber B, et al. What’s on the horizon for macroecology? Ecography. 2012 Aug 1;35(8):673–83.

29. Beck J, Ballesteros-Mejia L, Nagel P, Kitching IJ. Online solutions and the Wallacean shortfall what does GBIF contribute to our knowledge of species ranges? Divers Distrib. 2013;19(8):1043–1050.

30. Daru BH, Park DS, Primack RB, Willis CG, Barrington DS, Whitfeld TJS, et al. Widespread sampling biases in herbaria revealed from large-scale digitization. New Phytol. 2018 Jan 1;217(2):939–55.

31. Maldonado C, Molina CI, Zizka A, Persson C, Taylor CM, Alban J, et al. Estimating species diversity and distribution in the era of Big Data: to what extent can we trust public databases? Glob Ecol Biogeogr. 2015 Aug;24(8):973–84.

32. Meier R, Dikow T. Significance of Specimen Databases from Taxonomic Revisions for Estimating and Mapping the Global Species Diversity of Invertebrates and Repatriating Reliable Specimen Data. Conserv Biol. 2004 Apr 1;18(2):478–88.

33. Goodwin ZA, Harris DJ, Filer D, Wood JRI, Scotland RW. Widespread mistaken identity in tropical plant collections. Curr Biol CB. 2015 Nov 16;25(22):R1066–1067.

34. Zermoglio PF, Guralnick RP, Wieczorek JR. A Standardized Reference Data Set for Vertebrate Taxon Name Resolution. PLOS ONE. 2016 Jan 13;11(1):e0146894.

35. Wieczorek J, Guo Q, Hijmans R. The point-radius method for georeferencing locality descriptions and calculating associated uncertainty. Int J Geogr Inf Sci. 2004 Dec 1;18(8):745–67.

36. Dou L, Cao G, Morris PJ, Morris RA, Ludäscher B, Macklin JA, et al. Kurator: A Kepler package for data curation workflows. Procedia Comput Sci. 2012 Jan 1;9:1614–9.

37. Mathew C, Güntsch A, Obst M, Vicario S, Haines R, Williams A, et al. A semi-automated workflow for biodiversity data retrieval, cleaning, and quality control. Biodivers Data J. 2014;2:1–12.

38. Ponder W, Carter G, Flemons P, R. Chapman R. Evaluation of Museum Collection Data for Use in Biodiversity Assessment. Conserv Biol. 2001 Jun 1;15.

39. Boakes EH, McGowan PJ, Fuller RA, Chang-qing D, Clark NE, O’Connor K, et al. Distorted views of biodiversity: spatial and temporal bias in species occurrence data. PLOS Biol. 2010;8(6):e1000385.

40. Isaac NJ, Strien AJ, August TA, Zeeuw MP, Roy DB. Statistics for citizen science: extracting signals of change from noisy ecological data. Methods Ecol Evol. 2014;5(10):1052–1060.

41. Ruete A. Displaying bias in sampling effort of data accessed from biodiversity databases using ignorance maps. Biodivers Data J. 2015;(3):1–15.

42. Meyer C, Weigelt P, Kreft H. Multidimensional biases, gaps and uncertainties in global plant occurrence information. Ecol Lett. 2016 Aug 1;19(8):992–1006.

43. Meyer C, Jetz W, Guralnick RP, Fritz SA, Kreft H. Range geometry and socio-economics dominate species-level biases in occurrence information. Glob Ecol Biogeogr. 2016 Oct 1;25(10):1181–93.

44. Guralnick R, Van Cleve J. Strengths and weaknesses of museum and national survey data sets for predicting regional species richness: comparative and combined approaches. Divers Distrib. 2005 Jul 1;11(4):349–59.

45. Ball-Damerow JE, Oboyski PT, Resh VH. California dragonfly and damselfly (Odonata) database: temporal and spatial distribution of species records collected over the past century. ZooKeys. 2015;(482):67.

46. Rapacciuolo G, Ball-Damerow JE, Zeilinger AR, Resh VH. Detecting long-term occupancy changes in Californian odonates from natural history and citizen science records. Biodivers Conserv. 2017 Nov;26(12):2933–49.

47. Sierwald P, Bieler R, Shea EK, Rosenberg G. Mobilizing Mollusks: Status Update on Mollusk Collections in the U.S.A. and Canada. Am Malacol Bull. 2018 Dec;36(2):177–214.

48. ter Steege H, A. Persaud C. The phenology of Guyanese timber species—A compilation of a century of observations. Plant Ecol. 1991 Jan 9;95:177–98.

49. Peterson CH. Relative abundances of living and dead molluscs in two Californian lagoons. Lethaia. 1976 Apr 1;9(2):137–48.

50. Boag DA. Overcoming sampling bias in studies of terrestrial gastropods. Can J Zool. 1982 Jun;60(6):1289–92.

51. Dorazio RM. Accounting for imperfect detection and survey bias in statistical analysis of presence-only data. Glob Ecol Biogeogr. 2014 Dec 1;23(12):1472–84.

52. Zeilinger AR, Rapacciuolo G, Turek D, Oboyski PT, Almeida RPP, Roderick GK. Museum specimen data reveal emergence of a plant disease may be linked to increases in the insect vector population. Ecol Appl Publ Ecol Soc Am. 2017 Sep;27(6):1827–37.

53. Chapman AD. Uses of Primary Species-Occurrence Data, version 1.0. Report for the Global Biodiversity Information Facility. [Internet]. Copenhagen; 2005. Available from: Http://www.gbif.org/orc/?doc_id=1300.

54. Ariño A, Noesgaard D, Hjarding A, Schigel D. Biodiversity Information Services: A (not-so-) little knowledge that acts. Biodivers Inf Sci Stand. 2018 May 22;2:e25738.

55. Roy Rosenzweig Center for History and New Media. Zotero [Internet]. 2017. Available from: www.zotero.org/download

56. Ball-Damerow JE, Brenskelle L, Barve N, LaFrance R, Soltis PS, Sierwald P, et al. Bibliographic dataset characterizing studies that use online biodiversity databases [Internet]. Zenodo; 2019 [cited 2019 Mar 13]. Available from: https://zenodo.org/record/2589439#.XIlE5RNKjBI

57. Chavan V, Penev L. The data paper: a mechanism to incentivize data publishing in biodiversity science. BMC Bioinformatics. 2011;12(15):S2.

58. Moritz T, Krishnan S, Roberts D, Ingwersen P, Agosti D, Penev L, et al. Towards mainstreaming of biodiversity data publishing: recommendations of the GBIF Data Publishing Framework Task Group. BMC Bioinformatics. 2011;12(15):S1.

59. Whitlock MC. Data archiving in ecology and evolution: best practices. Trends Ecol Evol. 2011 Feb;26(2):61–5.

60. Smith V, Penev L. E-Infrastructures for Data Publishing in Biodiversity Science. PenSoft Publishers LTD; 2011. 425 p.

61. Costello MJ, Michener WK, Gahegan M, Zhang Z-Q, Bourne PE. Biodiversity data should be published, cited, and peer reviewed. Trends Ecol Evol. 2013 Aug;28(8):454–61.

62. Costello MJ, Wieczorek J. Best practice for biodiversity data management and publication. Biol Conserv. 2014 May 1;173:68–73.

63. Wilkinson MD, Dumontier M, Aalbersberg IjJ, Appleton G, Axton M, Baak A, et al. The FAIR Guiding Principles for scientific data management and stewardship. Sci Data. 2016 Mar 15;3:160018.

64. Mooney H, Newton M. The Anatomy of a Data Citation: Discovery, Reuse, and Credit. J Librariansh Sch Commun. 2012 May 15;1(1):eP1035.

65. Escribano N, Galicia D, Ariño AH. The tragedy of the biodiversity data commons: a data impediment creeping nigher? Database J Biol Databases Curation. 2018 Apr 9 [cited 2018 Dec 24];2018. Available from: https://www.ncbi.nlm.nih.gov/pmc/articles/PMC5892138/

66. Vines TH, Albert AYK, Andrew RL, Débarre F, Bock DG, Franklin MT, et al. The Availability of Research Data Declines Rapidly with Article Age. Curr Biol. 2014 Jan 6;24(1):94–7.

67. Klump J, Huber R. 20 Years of Persistent Identifiers – Which Systems are Here to Stay? Data Sci J. 2017 Mar 22;16(0):9.

68. McMurry JA, Juty N, Blomberg N, Burdett T, Conlin T, Conte N, et al. Identifiers for the 21st century: How to design, provision, and reuse persistent identifiers to maximize utility and impact of life science data. PLOS Biol. 2017 Jun 29;15(6):e2001414.

69. Stark PB. Before reproducibility must come preproducibility. Nature. 2018 May 24;557:613.

70. Cousijn H, Kenall A, Ganley E, Harrison M, Kernohan D, Lemberger T, et al. A data citation roadmap for scientific publishers. Sci Data. 2018 Nov 20;5:180259.

71. Force MM, Robinson NJ. Encouraging data citation and discovery with the Data Citation Index. J Comput Aided Mol Des. 2014 Oct;28(10):1043–8.

72. Costello MJ, Appeltans W, Bailly N, Berendsohn WG, de Jong Y, Edwards M, et al. Strategies for the sustainability of online open-access biodiversity databases. Biol Conserv. 2014;173:155–165.

73. Huang X, Hawkins BA, Qiao G. Biodiversity data sharing: Will peer-reviewed data papers work? BioScience. 2013;63(1):5–6.

74. Pimm SL, Alibhai S, Bergl R, Dehgan A, Giri C, Jewell Z, et al. Emerging Technologies to Conserve Biodiversity. Trends Ecol Evol. 2015 Nov 1;30(11):685–96.

75. Wood KR. Rediscovery, conservation status and taxonomic assessment of Melicope degeneri (Rutaceae), Kaua ‘i, Hawai ‘i. Endanger Species Res. 2011;14(1):61–68.

76. Costello MJ. Motivating Online Publication of Data. BioScience. 2009 May 1;59(5):418–27.

77. Costello MJ, Bouchet P, Boxshall G, Fauchald K, Gordon D, Hoeksema BW, et al. Global coordination and standardisation in marine biodiversity through the World Register of Marine Species (WoRMS) and related databases. PLOS ONE. 2013;8(1):e51629.

78. Tydecks L, Jeschke JM, Wolf M, Singer G, Tockner K. Spatial and topical imbalances in biodiversity research. PLOS ONE. 2018 Jul 5;13(7):e0199327.

79. Chapman AD. Numbers of Living Species in Australia and the World: A Report for the Australian Biological Resources Study [Internet]. Toowoomba, Australia: Australian Government Department of the Environment and Energy; 2009. Report No.: ISBN: 9780 642 56861 8. Available from: http://www.environment.gov.au/science/abrs/publications/other/numbers-living-species/contents#copyright

80. Sánchez-Fernández D, Lobo JM, Abellán P, Ribera I, Millán A. Bias in freshwater biodiversity sampling: the case of Iberian water beetles. Divers Distrib. 2008 Sep 1;14(5):754–62.

81. Ballesteros-Mejia L, Kitching IJ, Jetz W, Nagel P, Beck J. Mapping the biodiversity of tropical insects: species richness and inventory completeness of African sphingid moths. Glob Ecol Biogeogr. 2013 May 1;22(5):586–95.

82. Costello MJ, Wilson S, Houlding B. Predicting total global species richness using rates of species description and estimates of taxonomic effort. Syst Biol. 2012 Oct;61(5):871–83.

83. Rosenberg G. A New Critical Estimate of Named Species-Level Diversity of the Recent Mollusca*. Am Malacol Bull. 2014 Sep;32(2):308–22.

84. Schuh RT, Hewson-Smith S, Ascher JS. Specimen databases: A case study in entomology using web-based software. Am Entomol. 2010;56(4):206–216.

85. Mantle B, LaSalle J, Fisher N. Whole-drawer imaging for digital management and curation of a large entomological collection. ZooKeys. 2012 Jul 20;209:147–63.

86. Holovachov O, Zatushevsky A, Shydlovsky I. Whole-Drawer Imaging of Entomological Collections: Benefits, Limitations and Alternative Applications. J Conserv Mus Stud. 2014 Oct 29;12(1):Art. 9.

87. Hereld M, Ferrier NJ, Agarwal N, Sierwald P. Designing a High-Throughput Pipeline for Digitizing Pinned Insects. In: 2017 IEEE 13th International Conference on e-Science (e-Science). 2017. p. 542–50.

88. Price BW, Dupont S, Allan EL, Blagoderov V, Butcher AJ, Durrant J, et al. ALICE: Angled Label Image Capture and Extraction for high throughput insect specimen digitisation. 2018 Nov 5 [cited 2019 Mar 13]; Available from: https://osf.io/9p4f6/

89. Hoffmann M, Hilton-Taylor C, Angulo A, Böhm M, Brooks TM, Butchart SHM, et al. The Impact of Conservation on the Status of the World’s Vertebrates. Science. 2010 Dec 10;330(6010):1503–9.

90. Pino-del-Carpio A, Ariño AH, Miranda R. Data exchange gaps in knowledge of biodiversity: implications for the management and conservation of Biosphere Reserves. Biodivers Conserv. 2014;23(9):2239–2258.

91. Pino-Del-Carpio A, Villarroya A, Ariño AH, Puig J, Miranda R. Communication gaps in knowledge of freshwater fish biodiversity: implications for the management and conservation of Mexican biosphere reserves. J Fish Biol. 2011 Dec;79(6):1563–91.

92. Ball J, Beche L, Mendez P, H. Resh V. Biodiversity in Mediterranean-climate streams of California. Hydrobiologia. 2013 Nov 1;719.

93. Dewalt E, Favret C, W. Webb D. Just how imperiled are aquatic insects? A case study of stoneflies (Plecoptera) in Illinois. Ann Entomol Soc Am. 2005 Oct 31;98:941–50.

94. Ball-Damerow JE, M’Gonigle LK, Resh VH. Changes in occurrence, richness, and biological traits of dragonflies and damselflies (Odonata) in California and Nevada over the past century. Biodivers Conserv. 2014 Jul 1;23(8):2107–26.

95. Colla SR, Gadallah F, Richardson L, Wagner D, Gall L. Assessing declines of North American bumble bees (Bombus spp.) using museum specimens. Biodivers Conserv. 2012;21(14):3585–3595.

96. Hallmann CA, Sorg M, Jongejans E, Siepel H, Hofland N, Schwan H, et al. More than 75 percent decline over 27 years in total flying insect biomass in protected areas. PLOS ONE. 2017 Oct 18;12(10):e0185809.

97. Escribano N, Ariño AH, Galicia D. Biodiversity data obsolescence and land uses changes. PeerJ. 2016;4:1–15.

98. Peterson AT, Soberón J, Krishtalka L. A global perspective on decadal challenges and priorities in biodiversity informatics. BMC Ecol. 2015;15(1):15.

99. Austin M, Van Niel K. Improving species distribution models for climate change studies: Variable selection and scale. J Biogeogr. 2010 Nov 9;38:1–8.

100. Stanton JC, Pearson RG, Horning N, Ersts P, Reşit Akçakaya H. Combining static and dynamic variables in species distribution models under climate change. Methods Ecol Evol. 2012;3(2):349–357.

101. Fournier A, Barbet-Massin M, Rome Q, Courchamp F. Predicting species distribution combining multi-scale drivers. Glob Ecol Conserv. 2017 Oct 1;12:215–26.

102. Staniczenko PPA, Sivasubramaniam P, Suttle KB, Pearson RG. Linking macroecology and community ecology: refining predictions of species distributions using biotic interaction networks. Ecol Lett. 2017 Jun 1;20(6):693–707.

103. Ratnasingham S, Hebert PDN. bold: The Barcode of Life Data System (http://www.barcodinglife.org). Mol Ecol Notes. 2007 May 1;7(3):355–64.

104. Bold Systems v4 [Internet]. [cited 2019 Apr 5]. Available from: http://www.boldsystems.org/

105. Hinchliff CE, Smith SA, Allman JF, Burleigh JG, Chaudhary R, Coghill LM, et al. Synthesis of phylogeny and taxonomy into a comprehensive tree of life. Proc Natl Acad Sci. 2015 Oct 13;112(41):12764–9.

106. Chavan V, Berents P, Hamer M. Towards demand driven publishing: approaches to the prioritisation of digitisation of natural history collections data. Biodivers Inform [Internet]. 2010 Oct 9 [cited 2016 Aug 23];7(2). Available from: https://journals.ku.edu/index.php/jbi/article/view/3990

107. Rios, N. E., Bart, HL. GEOLocate. Belle Chasse, LA: Tulane University Museum of Natural History. Available from: http://www.geo-locate.org

108. Boyle B, Hopkins N, Lu Z, Raygoza Garay JA, Mozzherin D, Rees T, et al. The taxonomic name resolution service: an online tool for automated standardization of plant names. BMC Bioinformatics. 2013;14(1):16.

109. Chamberlain SA, Szöcs E. taxize: taxonomic search and retrieval in R. F1000Research [Internet]. 2013 Oct 28 [cited 2018 Oct 10];2. Available from: https://www.ncbi.nlm.nih.gov/pmc/articles/PMC3901538/

110. Froese R, Pauly D. FishBase. World Wide Web electronic publication. 2014 Jan 2 [cited 2019 Mar 27]; Available from: https://www.scienceopen.com/document?vid=dc419213-0ca3-48cc-901c-2934ecf4441e

111. WoRMS Editorial Board. World Register of Marine Species. Available from http://www.marinespecies.org at VLIZ. Accessed yyyy-mm-dd. [Internet]. VLIZ; 2017 [cited 2019 Apr 5]. Available from: http://www.marinespecies.org/imis.php?dasid=1447&doiid=170

112. MilliBase [Internet]. [cited 2019 Apr 2]. Available from: http://www.millibase.org/

113. MolluscaBase - Introduction [Internet]. [cited 2019 Apr 2]. Available from: http://www.molluscabase.org/

114. Ball-Damerow JE, Mendez PK, Sierwald P, Bieler R, Yoder M, DeWalt RE. Taxonomic data quality in GBIF: a case study of aquatic macroinvertebrate groups. In Ann Arbor, MI; 2017.

115. Wägele H, Klussmann-Kolb A, Kuhlmann M, Haszprunar G, Lindberg D, Koch A, et al. The taxonomist - an endangered race. A practical proposal for its survival. Front Zool. 2011 Oct 26;8:25.

116. Drew LW. Are We Losing the Science of Taxonomy?: As need grows, numbers and training are failing to keep up. BioScience. 2011 Dec;61(12):942–6.

117. Vaidya G, Lepage D, Guralnick R. The tempo and mode of the taxonomic correction process: How taxonomists have corrected and recorrected North American bird species over the last 127 years. PLoS ONE [Internet]. 2018 Apr 19 [cited 2019 Mar 27];13(4). Available from: https://www.ncbi.nlm.nih.gov/pmc/articles/PMC5909608/

118. Arvanitidis CD, Warwick RM, Somerfield PJ, Pavloudi C, Pafilis E, Oulas A, et al. Research Infrastructures offer capacity to address scientific questions never attempted before: Are all taxa equal? PeerJ Inc.; 2018 Aug [cited 2019 Mar 27]. Report No.: e26819v2. Available from: https://peerj.com/preprints/26819

119. Otegui J, Guralnick RP. The geospatial data quality REST API for primary biodiversity data. Bioinformatics. 2016;32(11):1755–1757.

120. Paul D, Fisher N. Challenges For Implementing Collections Data Quality Feedback: synthesizing the community experience. Biodivers Inf Sci Stand. 2018 Jun 13; 2:e26003.

